# Sound preferences in mice are sex-dependent

**DOI:** 10.1101/2024.08.04.606510

**Authors:** Kamini Sehrawat, Israel Nelken

## Abstract

We investigated the impact of early exposure to sound and to silence on sound preferences later in life in mice. We exposed young mice during the critical periods to excerpts of music (first movement of Beethoven’s symphony no. 9), non-music sounds, or to silence. We tested the sound preference behavior a few weeks later. Music exposure affected mouse behavior in a sex-dependent manner: male mice largely preferred the environment to which they were exposed, while female mice showed a weak reduction in their seemingly inborn aversion to sound. The neural activity in auditory cortex was suppressed in exposed compared to naive mice, regardless of exposure type. Remarkably, a robust negative correlation was found between neural response and behavior in female, but not in male, mice.

## Introduction

Sounds often have emotional content, irrespective of their importance for guiding behavior. In humans, some sounds are preferred (1) while others evoke strong aversive reactions (2, 3). Sound preferences may be innate, as is the attraction of infants to speech sounds (4), the innate caregiving tendencies driven by infant sounds observed in adult humans (5), or the preference of rats for the conspecific 50-kHz calls (‘laugh calls’), which are linked to positive behaviors such as mating and playing (6).

In humans, sound preferences are mostly tested using music. Music preferences are likely learned, rather than innate (7–9). They have been linked to the activation of emotionally relevant brain regions such as nucleus accumbens (NAcc) (1, 10, 11). Pleasurable music is associated with the release of dopamine in NAcc (12, 13), presumably from the ventral tegmental area (VTA) (14). Notably, VTA activity exhibited a significant correlation with NAcc activity, implying an interplay between these regions in modulating responses to pleasant music (15). Moreover, the functional connectivity between NAcc and auditory cortex increased for preferred music, and was reduced in individuals with music anhedonia (16, 17). This body of work suggests that preferred music causes heightened interactions between the auditory cortex (AC), NAcc, and VTA, and the resulting neural activity (including dopamine release) underlies music preferences.

We wanted to study sound preferences in animal models. For that purpose, we took advantage of the finding that sound preferences in adult mice can be influenced by early life experiences, particularly during critical periods. Critical periods are specific time windows in early stages of postnatal development, characterized by high level of brain plasticity in response to sensory experiences (18). Auditory experience during critical periods causes long term alterations in the circuitry of auditory cortex (19–21) as well as changes in behavior. For example, adult female zebra finches developed preferences for songs they have been exposed to during their rearing period (22). Importantly, studies on mice have shown that exposure to specific sound environments during the critical periods lead to a preference for those auditory environments in adulthood (23, 24).

Here, we study sound preferences in mice induced by early exposure (P7-P40) to music, and related them to auditory-evoked activity in the auditory cortex. Greatly extending previous reports (23, 24), we found that the effects were strongly sex-dependent: sound exposure affected consistently the behavior of male, but much less so that of female, mice; and neural activity in auditory cortex was negatively correlated to music preference in female, but not in male, mice.

## Results

### Sex dependent preference for music in mice

We exposed C57BL/6 mice of both sexes to excerpts from the first movement Beethoven’s Symphony No. 9 or to silence during their infancy. Exposure was conducted in the home cage, with free access to food and water (P7-P40, (21); Fig. 1A). Exposure was intermittent and mild.

**Fig. 1.**
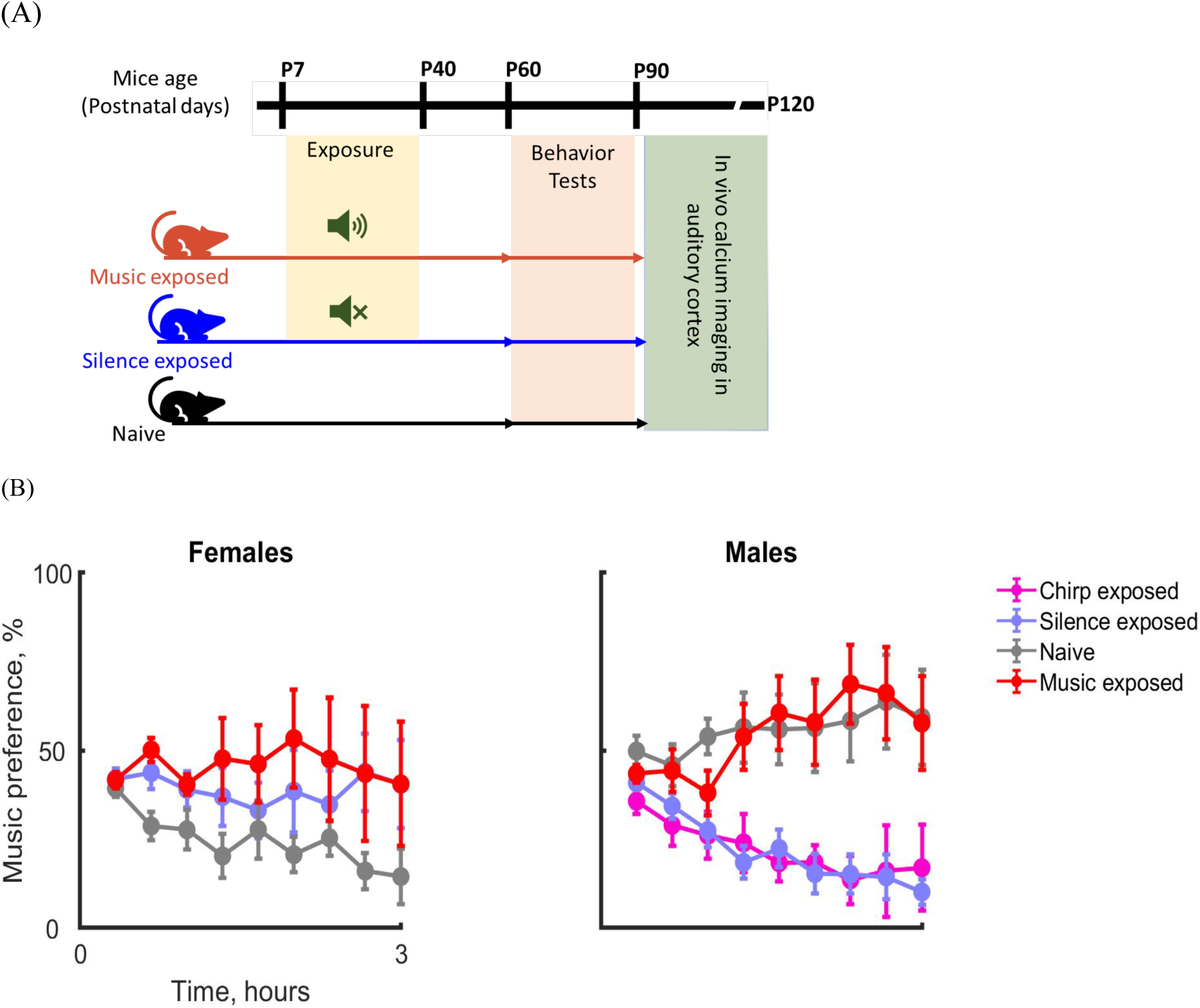
Sex dependent preference for music in mice. **(A**) Schematic of the experimental methods. Music or silence exposure was conducted between P7-P40. Behavioral tests were performed between P60-P90, followed by the acute widefield calcium imaging in auditory cortex. **(B)** Time spent in the music zone (Music preference, %), for the males (left) and females (right) in all groups (mean ± SE; red: Music-exposed, M=11, F=6; blue: Silence-exposed, M=10, F=8; black: Naïve, M=10, F=6); magenta: Chirp-exposed, M=7, F=0).

Silence exposure consisted of daily 6 hour sessions in a sound-proof chamber, and music exposure consisted of similar sessions, during which the sounds were presented for 4 periods of 20 minutes with 60 minutes break between them. A group of naïve mice, that remained in the animal facility from birth and until the behavioral test, served as an additional control.

When the mice reached adulthood (>P60), they were tested for sound preferences. The mice explored a box for 180 minutes (without reinforcement). Half of the box was used as a silent zone and half as a music zone. Mice tended to reduce their activity level over time (Fig. S1; Table S1), sometimes settling in one or the other side of the box. Music preference was quantified by the amount of time spent in the music zone in nine time bins, each lasting 20 minutes. There was a significant effect of group (Table S2; here and elsewhere, permutation test; main effect of group, P=0.016). In fact, the silence exposed animals spent overall less time in the music zone than naïve animals (post-hoc test, P<0.001) while the music exposed animals spent overall more time in the music zone than naïve animals (post-hoc test, P<0.001).

Unexpectedly, male and female mice showed different behavioral patterns in this test (Figs. 1B, S2; Table S2, main effect of sex, P=0.021 and group X sex interaction, P=0.007). Naïve male and female mice already showed difference in their patterns of preference (post-hoc test, P=0.003): the naïve females strongly avoided the music zone while the males showed a mixed pattern, with some of the naïve males settling in the music zone and some in the silence zone. The effect of exposure on male mice was large and robust (Table S3). Male mice exposed to silence in infancy showed a strong preference for the silent zone, while mice exposed to music in infancy showed a mixed pattern of preferences similar to that of the naïve mice. On the other hand, female mice exposed to silence or to music in infancy spent on average somewhat longer periods of time in the music zone relative to naïve mice. However, all groups of female mice had substantial heterogeneity and these differences did not reach significance (Table S4).

Finally, the effect of music exposure on the male mice was specific to the exposed music: males exposed to chirps, rather than to the music, avoided the music zone during the test to a degree similar to that of the silence exposed males (Table S5).

### Suppression of neural responses in exposed mice

After completing the behavioral tests, we measured sound responses under anesthesia in the same mice using wide-field calcium imaging. We used broadband noise (BBN), pure tones, music excerpts from the sound file used in the behavioral experiments (familiar sounds; Fig S3A), as well as control complex sounds (unfamiliar sounds; Fig S3A).

Remarkably, most exposed mice exhibited a significant suppression of the peak calcium responses compared to naive animals (Fig. 2A and S3B; Table S6; main effect of group, P=0.035; post-hoc permutation test for the contrast (naive-(music-exposed+silence-exposed)/2), P=0.009). There was a significant main effect of sex (P=0.048) and the significant interaction of sex with the stimulus class (P=0.025). Given this significant interaction between the sex and stimulus class we analyzed the three stimulus class (BBN, pure tone, and familiar sounds) separately. And, since our experimental design included a specific comparison between the silence- and the music-exposed groups, we further studied the peak responses of these two groups, omitting the naïve group. The peak response to pure tones (P=0.036; Table S8) and familiar sounds (P=0.002; Table S7) varied between music and silence exposed groups. The responses to the familiar sounds had a highly significant interaction between group and sex (P=0.029): they were larger in music-exposed than in silence-exposed females (Fig. 2B bottom, Table S7), but smaller in music-exposed than in silence-exposed males. On the other hand, the responses to pure tones showed a reverse pattern (Fig. 2B, middle; Table S8). The responses to broadband noise didn’t show a significant effect of exposure group or sex (Fig. 2B, top; Table S9). Remarkably, the responses in chirp-exposed males were similar to those of the silence-exposed males and differed significantly from those of the music-exposed males (Fig. S3B; Table S10).

**Fig. 2.**
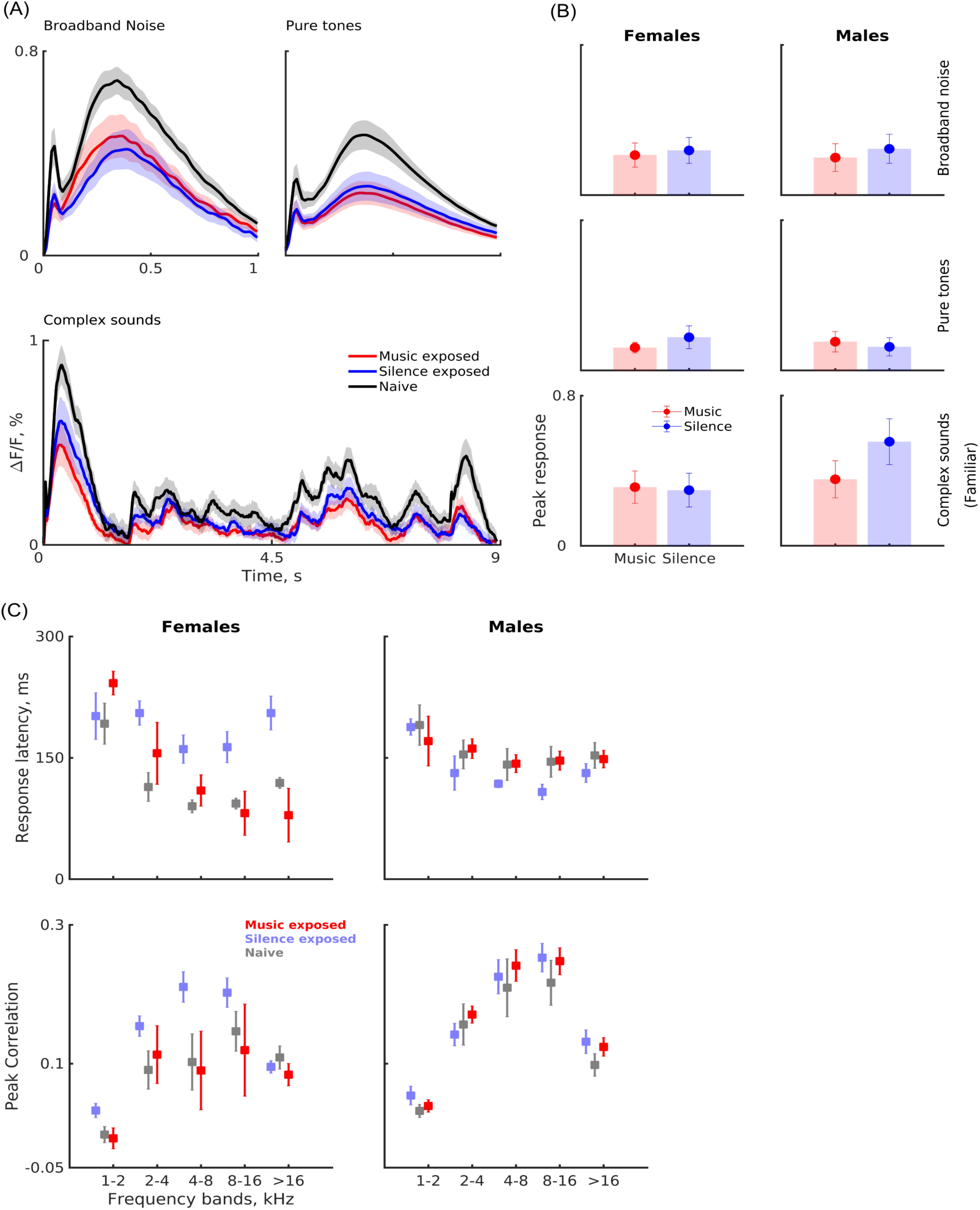
Effects of early exposure on neuronal responses in auditory cortex **(A**) Average neural responses (mean ± SE) as a function of time, for all sound types: Broadband noise (top left; red: music-exposed; blue: silence-exposed; black: naïve), Pure tones (top right), and familiar complex sounds (bottom). **(B)** Peak responses (mean ± SE) of music and silence exposed males (right) and females (left) for broadband noise (top), Pure tones (middle) and familiar sounds (bottom). **(C)** Response latencies (top) and peak correlations (bottom) of the neural responses with sound envelopes in different frequency bands for familiar sounds, in females (left; red: music-exposed; blue: silence-exposed; black: naïve) and males (right).

For the complex sounds, we also studied the moment-by-moment relations between sound waveform and neuronal responses, by correlating the calcium waveforms with the output of octave-wide frequency bands that averaged the simulated responses of the auditory periphery (see supplementary methods; Fig. S4). For the familiar sounds, the latency of the peak correlation showed a highly significant group X sex interaction (P=0.003; Fig. 2C; Table S11): in female mice, but not in male mice, the latency of the peak correlations (reflecting the duration of the firing events that lead to calcium entry following an excitatory acoustic event) was significantly longer in silence-exposed animals than in music-exposed animals (and in naïve animals). Peak correlations also showed a significant band X group X sex interaction (P=0.031; Fig. 2C; Tables S12). For unfamiliar sounds, most effects were non-significant, with some weak interactions (Fig. S5; Tables S13, S14).

### Neural correlates of preference behavior

We compared the peak calcium responses analyzed in Fig. 2 with the behavior of the mice analyzed in Fig. 1, on an animal-by-animal basis (Fig. 3). The relationship between the two was significantly different in males and females (group X sex interactions were significant for all sound types; Table S15). In male mice, the correlations between response magnitude and time in the music zone were not significant for any sound type (Fig. 3, Fig. S6, Tables S16, S17A). On the other hand, for females, the correlations between response magnitude and time in the music zone were large and significant for all sound types (pure tones: P=0.028, broadband noise: P=0.030, all complex sounds: P=0.002; for the subset of familiar sounds: P=0.004). Remarkably, these correlations were negative, so that activity in auditory cortex was aversive for female mice.

**Fig. 3.**
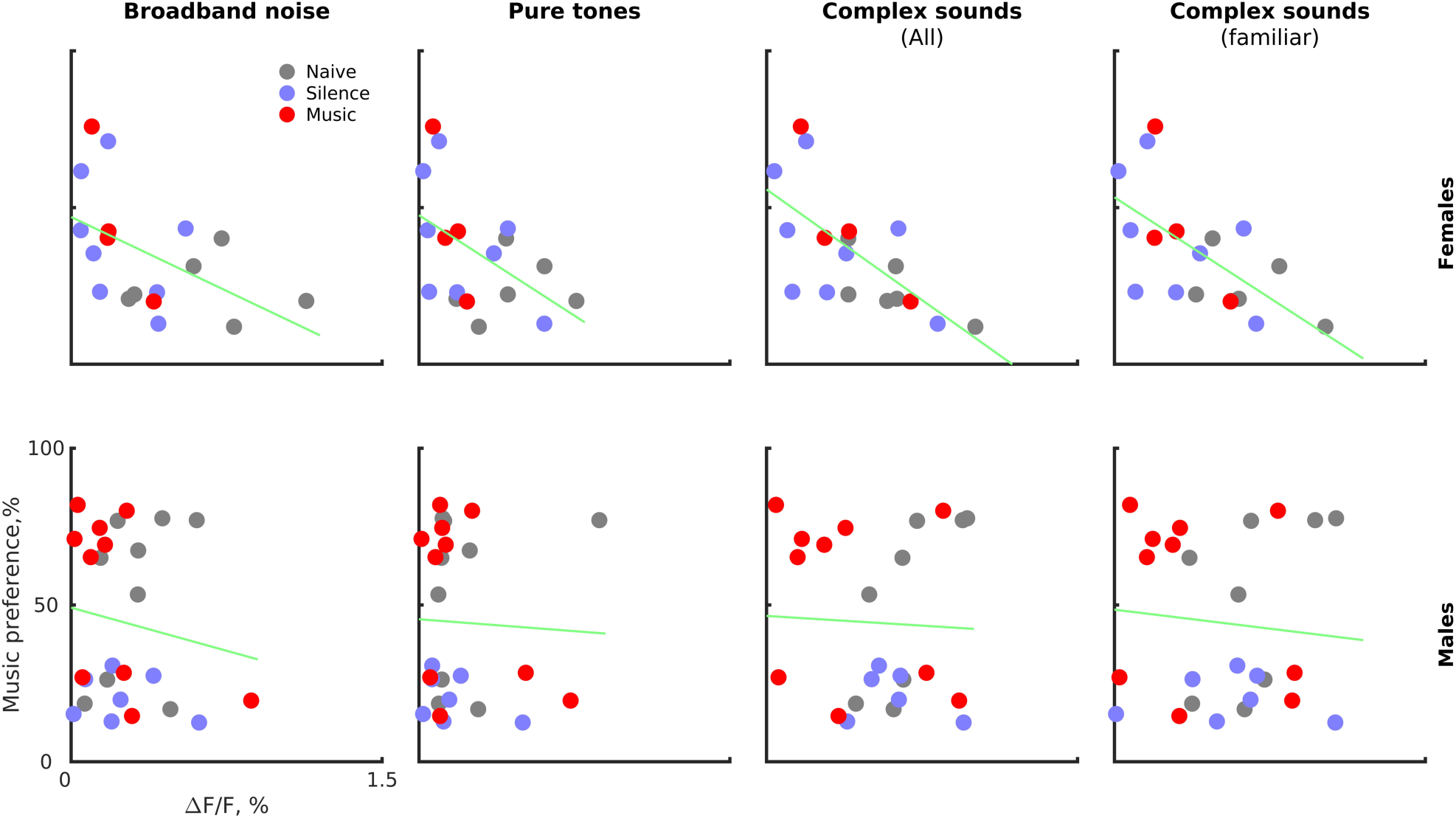
**Neural correlates of preference behavior**. Peak neural responses (x-axis) plotted against music preference of the same animal (y-axis) for females (top) and males (bottom). The correlations are sex-dependent, but within sex, they are consistent for all sound types

## Discussion

We investigated the influence of sound exposure during the critical periods of development of the mouse auditory system on sound preference behavior later in life. The exposure period we used encompassed both early and late critical periods (21). Following a modified version of previous exposure protocols (23, 24), we exposed the mice to excerpts from the first movement of Beethoven’s 9th symphony during days P7-P40. We extended previous results by adding control groups consisting of chirp-exposed animals and of naïve animals; by recording neural activity from the auditory cortex of these mice; and by correlating neural activity in auditory cortex with behavior.

### Behavior

Our findings reproduced previously reported behavioral consequences of early exposure to music in adult male mice. Notably, in male mice, we observed a clear tendency for music-exposed mice to spend more time in the music zone relative to the silence-exposed mice, as in Yang et al. (2012). Silence-exposed male mice consistently preferred the silence zone. Music-exposed males exhibited a bimodal pattern of preference, with some clearly favoring the music zone, while others preferred the silence zone. The behavioral effects were specific to the sounds to which the mice were exposed: the chirp-exposed males preferred the silence zone over the music zone. Similar findings have been used to argue that early exposure to sounds leads to the development of preference to the exposed sounds in mice (23, 24).

However, our results show that such conclusions are premature. First, by testing a group of naïve mice who were not exposed to either music and silence, we discovered that it was not music exposure that modified mouse behavior – naïve and music-exposed mice spent on average about the same amount of time in the music zone. Instead, it was exposure to other environments – silence or chirps – that caused male mice to avoid the music zone later in life, producing the robust difference between music-exposed and silence- or chirp-exposed male mice. We conclude that while early exposure to sound environments has significant effect on male mouse behavior, this effect is largely negative – exposed male mice avoided novel sound environments, but did not prefer familiar ones more than naïve mice do (Fig. 1B left, Fig. S2 bottom).

Second, the consequences of early exposure were sex dependent. Female mice were much less affected by early exposure than male mice: they showed a somewhat increased tendency to spend time in the music zone following either music- or silence-exposure, although the increase was not statistically significant.

The estrous cycle could underlie the larger behavioral variability we observed in females. However, previous research has shown that in female mice, the impact of hormonal cycle on exploratory behavior is small (25). Therefore, we believe that it was not the origin of the sex differences we observed. Sex-dimorphic behaviors, evident in mating or aggression, evolved from the interplay between genetics and neural circuitry across sexually reproducing species (26–28). Sound preferences as tested here may be one of these sex-dimorphic behaviors.

### Neural activity

Generally, early exposure to pure tones and other narrowband stimuli seems to cause an increase in the responses to the exposed sounds later in life. Thus, when rat pups were exposed to pulsed tones during the early critical period of development, there was a selective enlargement of the cortical representations corresponding to the frequency utilized in the exposure stimuli accompanied by disrupted tonotopicity and wider receptive fields (29, 30). On the other hand, early exposure to broadband stimuli may result in suppressed responses in auditory cortex. Thus, evoked activity in auditory cortex has been suppressed in rats reared in a complex acoustic environment with a positive reinforcement (31). Rats exposed to pulsed noise during both early and late critical periods showed reduced neural responses to pure tones as compared to a control naïve group (32). Such suppression has been reported in other species as well: exposure to multitone sounds in juvenile cats led to suppressed responses to sounds in primary auditory cortex (33).

In our experiments, exposure to music, which is complex and broadband, resulted in suppression of the activity in auditory cortex as compared to naïve mice. This result is consistent with previous studies that used broadband stimuli. However, the silence-exposed mice in our study also had reduced neural activity compared to naïve controls. In some studies, differences were reported in the neural responses between mice reared in silence environment compared to mice reared in normal housing (34), while in other studies, such differences were not apparent (29). Silence-exposure in our study differed from that used in previous studies: the mice spent most of the day in the animal facility, exposed to ambient sounds, and were exposed to silence only intermittently. One way to interpret our findings is to consider that both music- and silence- exposed mice alternated between multiple sound environments with different properties – music, silence, and normal housing sounds in one case: silence and normal housing sounds in the other. It may be that such alternations led to the suppression of the activity in auditory cortex.

### Relationships between behavior and activity in auditory cortex

In humans, preferred sounds are associated with increased functional correlation between activity in auditory cortex and in ventral striatum, as well as increased dopamine release. Translated to the mouse model, we expected to see an increased response to preferred sounds in auditory cortex as well as a positive correlation between sound-evoked activity and behavioral music preference. Instead, we discovered a subtler picture.

Our experimental design made it possible to compare the activity in auditory cortex and music preferences in the same mice (admittedly, recorded under anesthesia). In males, the correlation between auditory cortex activity and behavior was small and not significant (Fig. 3 bottom). In contrast, in female mice, the correlation between auditory cortex activity and behavior was highly significant for all sound types (Fig. 3 top). Surprisingly, the correlation was negative, suggesting that female mice preferred low levels of auditory cortex activity – auditory cortex activity was aversive.

To account for these results, we hypothesize that as in humans, preferred sounds increase the activity of dopaminergic (DA) neurons in the VTA, leading to the observed behavioral preferences. We are unaware of any direct projection from the auditory cortex to the VTA. On the other hand, the VTA receives auditory inputs indirectly through both cortical and subcortical secondary targets of the auditory system (35–37).

We therefore posit that the auditory cortex influences activity in the VTA through an intermediate station that serves as a gate which determines sound preferences. In females, this gate remains predominantly open, transmitting auditory system activity to the VTA. Since larger responses in the auditory cortex of female mice correlate with shorter dwell times in the music zone (Fig. 3, top), the model suggests a net inhibitory effect of auditory activity on DA neurons in female mice, potentially by a projection that terminates on the GABAergic neurons of the VTA (38). In contrast, in males, the lack of correlation between the preference behavior and sound driven responses in auditory cortex indicates that the gate is closed to auditory cortex responses. Instead, auditory effects in the male VTA may be mostly driven by subcortical inputs.

We show here that the effects of early exposure to sound environments have substantial consequences on behavior and on neural activity in the adult brain, but that these effects are sex dependent. Thus, sound preferences may be driven by different mechanisms in male and female mice.

## Materials and Methods

### Animals

The experimental procedures were approved by the Institutional Animal Care and Use Committee at the Hebrew University of Jerusalem, Israel. The Hebrew University is an AAALAC approved institution. C57BL/6J mice, including both males and females, were used in the study. Pregnant female mice were housed in pairs with adequate bedding and free access to food and water under a 12:12-hour light/dark cycle. The pups were kept with their mothers until P20, and at P24 the mice were sex-separated and housed with their same-sex littermates. The mice had free access to food and water throughout.

### Exposure paradigm

Four groups of mice were used in these experiments: naïve (n=16; 10 males, 6 females), exposed to music (n=17; 11 males, 6 females), exposed to chirps (n=8; 7 males, 1 females), and exposed to silence (n=18; 10 males, 8 females). The naive mice were housed in the animal facility and handled in a similar manner to the other groups but were not brought to the lab for the exposure sessions. The exposure sessions were conducted between postnatal day 7 to 40 (including weekends) to cover both early and late critical periods (21) . The exposure sessions of the music-exposed group took place during one half of the day and those of the silence-exposed group during the other half. These half-days were switched between the two groups every other day, so that both music- and silence-exposed mice was exposed in the mornings as well as in the afternoons. The exposed groups were brought from the animal facility to the lab and placed in an acoustic chamber in their home cage with unrestricted access to water and food. The music used for exposure was a 5-minute excerpt from the first movement of Beethoven’s Symphony No. 9, played in a repetitive loop for 20 minutes. The sound was high-pass filtered (corner frequency = 1500 Hz) to remove low frequency components that are inaudible to the mice, and then presented at a sound level of 70-75 dB SPL. The 20-minute music exposure was repeated four times every day, with a silent interval of 60 minutes between exposures. For the silence-exposed group, the same procedure was followed except that the speakers were not activated. The chirp-exposed group was tested separately using the same procedure. Upward and downward frequency-modulated sweeps covered the frequency range of 250 Hz to 50 kHz. The frequency changed linearly on a logarithmic frequency axis, and each chirp had a duration of 100 ms with 5 ms linear onset and offset ramps. A chirp sequence consisted of a pair of downward and upward chirps (each 100 ms long) separated by an interval of 400 ms, followed by 2400 ms of silence. These 3-second chirp sequences were looped for a total duration of 20 minutes, also presented at a sound level of 70 to 75 dB SPL. Prior to each experiment, the sounds were calibrated in situ (microphone: Brüel & Kjær Type 4189-A-021, Denmark).

### Behavioral experiments

All behavioral tests were conducted between postnatal days P60 and P90. The setup for assessing sound preferences in mice was adapted from previous studies (23, 24). We used a cubic box measuring 45 cm X 45 cm X 45 cm, placed within an acoustic chamber to ensure noise isolation. The box had two corners with red shelters and bedding material for nesting. The mouse location was tracked using a video camera (Ethovision-XT, Noldus, Wageningen, Netherlands) in ambient light. Two speakers (MF1, Tucker Davis Technology, USA) were installed, one above each shelter corner. The experimental box was divided by an unmarked diagonal boundary into two zones, a music zone and a silence zone, with one shelter in each zone. The software triggered the sound from the speaker located at the designated music zone when the mouse crossed the boundary between the zones. The location of the mouse was determined at a rate of 25/s. Each mouse spent a 90-minute habituation period in the test box, during which no auditory stimuli were presented. On the following day, the main three-hour music vs. silence test was conducted. The side of the music zone was determined as the zone in which the mouse spent the lesser amount of time during the habituation period on the previous day. During the test, the speaker located on top of the music corner emitted music when the animal was in the music zone of the box. The music played in this test consisted of the file from Beethoven 9th symphony used during the exposure of the music group, with the same filtering and sound level as during exposure. This sound file was used for testing all mice (naïve, exposed to music, to silence, and to chirps). Following the test of each mouse, thorough cleaning of the behavioral apparatus was performed using a solution of virosol (5%) and alcohol (70%).

### In-vivo widefield calcium imaging

The mice were initially anesthetized using sevoflurane at a concentration of 4% in 0.1 l/m O2 inside a custom-made closed box. Once profoundly anesthetized, the animal was carefully moved to the surgery table. A custom-made face mask was used to respirate the animal with a mixture of O2 and sevoflurane (EZ-155SF Stand-Alone Vaporizer, USA). A heating pad with a DC temperature controller (FHC 40-90-8D) was used to maintain the animal’s rectal temperature at 36±1℃. To protect the eyes, synthomycin 3% ointment (chloramphenicol 3%, Teva pharmaceutical industries Ltd., Netanya, IL) was applied. During the surgery, the depth of anesthesia was continuously monitored by assessing the rate and pattern of breathing using a strain gauge (KFH-06-120-C1-11L1M2R; Omega Engineering Inc, USA) positioned underneath the animal. The electrical signal from the breathing strain gauge was filtered between 3 and 30 Hz, amplified (gain = 10), and monitored online. The depth of anesthesia was confirmed by the absence of tail pinch and eyelid reflexes and by a respiration rate between 80 and 120 breaths per minute. Prior to the surgical procedure, 1μl of lidocaine 0.2% solution (Esracain 2%, RAFA laboratories, Jerusalem, IL) was injected subcutaneous under the scalp. After 10-15 minutes, the skin covering the dorsal surface of the skull was removed using fine scissors. The skull was then scraped using a surgical scalpel and dried with air flow. A custom-made metal bar was affixed to the midline of the dry skull using dental cement (Coral fix) mixed with 2-3 drops of histoacryl (B. Braun surgical S.A.). The cement hardened within 10-15 minutes, allowing the metal bar to be securely inserted into the holder. To access the temporal bone, the animal’s head was rotated 30-40 degrees using the metal bar holder. The skin over the temporal bone was incised, and the underlying muscles were scraped down with a scalpel to expose the skull surface. Metal retractors were used to maintain the position of these muscles throughout the procedure. The exposed area was then cleaned with saline solution and dried. A dental cement well was formed around the designated craniotomy site. A biopsy punch with a diameter of 3mm (Integra Miltex) was used to mark the location of the craniotomy. The craniotomy itself was performed using a micromotor drill (Microomotor schick Q profi, South Africa; Integra Miltex DHP1 carbide burs, Integra Lifesciences, USA). Throughout the procedure, the brain was kept moist with saline solution (0.9% w/v sodium chloride, Mini-Plasco, B. Braun).

After the surgery, the mouse was transferred from the surgical table to the calcium imaging setup. Acetoxymethyl (AM) ester form of the calcium-sensitive dye Oregon Green BAPTA-1 (OGB-1 AM, Molecular Probes, Eugene, OR, USA; 50 micrograms) was dissolved in 4 µl of dimethyl sulfoxide (DMSO) containing 20% Pluronic F-127 acid (Invitrogen by Thermo Fisher Scientific, USA). The dye solution was diluted in an extracellular buffer consisting of 150 mM NaCl, 2.5 mM KCl, and 10 mM HEPES to a final concentration of 0.5 mM. The dye solution was subjected to 1 minute of sonication, followed by filtration using a 0.2-micron syringe filter (PALL corporation nanosep, England, UK) to eliminate dye aggregation. A glass pipette (Borosilicate glass with filament, OD: 1.5 mm, ID: 0.86 mm, BF150-86-10; Shutter Instrument, Novato, California, USA) with a resistance of 3-4 MΩ was filled with 5-7 µl of the calcium dye solution. The pipette was positioned at an angle of 45 degrees relative to the sagittal midline and lowered to a depth of 300 μm from the surface of the cortex. A negative pressure of 3 PSI was applied to the pipette using a pressure pump (Pneumatic Picopump PV830, World precision instruments, USA). Once the pipette reached the desired depth, a pressure of 10 PSI was applied for 1 minute. These parameters have been previously determined for staining a sphere with a diameter of 400- 800 µm. To cover the entire craniotomy area of the auditory cortex, 4-6 injections were performed. Following the staining procedure, a glass window with a diameter of 3 mm was securely affixed over the craniotomy site for wide-field imaging. Recordings started 45-60 minutes after the end of the injections, to ensure sufficient staining of the neural tissue. During the recordings, the concentration of sevoflurane was lowered to 1.5-2% in 0.1 l/m O2. The recordings were conducted within a sound-proof chamber (IAC1202). The imaging setup consisted of a macroscope (Axio Zoom V16, Zeiss) with a 25 mm field of view objective lens (EC Plan-Neofluar 1x/0.025). Frame acquisition was performed by an sCMOS camera (ZYLA 4.2 PLUS, 4.2 Megapixel, Andor, Oxford instruments, UK) mounted on the macroscope. The excitation light was generated by Compact Light Source HXP 120 V (Leistungselektronik JENA GmbH). The field of view was adjusted to image the craniotomy. The imaging data were acquired using the Andor Solis software (Andor, Oxford Instruments, UK). The images were binned to 512 x 512 pixels and stored for offline analysis (average magnification: 39X, average pixel size: 15.2×15.2 μm^2^).

### Acoustic stimuli

Simple sounds, such as pure tones and broadband noise (BBN), were generated online at a sampling rate of 192 kHz using MATLAB (MathWorks). The sounds were transduced into voltage signals using a sound card (RME HDSP9632) and attenuated by 3 dB using an analog attenuator (PA5, TDT). The attenuated signals were presented using a speaker (MF1, TDT) positioned near the right ear of the mouse. During sound calibration, we determined that an attenuation level of 0 dB corresponded to sound pressure levels (SPL) ranging from 90-100 dB for pure tones up to 40 kHz and 80-90 dB SPL for frequencies between 40-50 kHz. BBN bursts were generated at a spectrum level of -60 dB/Hz relative to the pure tones. BBN bursts had a duration of 100 ms, with linear onset and offset ramps of 10 ms and an inter-stimulus interval (ISI; onset to onset) of 1000 ms. Pure tone stimuli were presented in quasi-random frequency sequences consisting of 190 pure tone bursts of 19 frequencies, ranging from 1 kHz to 64 kHz (3 frequencies/octave, equally spaced on a logarithmic frequency axis). Each frequency was repeated 10 times within the sequence. Each pure tone burst had a duration of 100 ms and a rise/fall time of 5 ms and the ISI was 1000 ms.

Complex sounds consisted of 8 second excerpts (n = 3) from the first movement of Beethoven’s Symphony No. 9, extracted from the same file used for exposure and behavioral testing; music excerpts (n = 3) from classical Indian ragas (Mishra Khamaj; by Pandit Shiv Kumar Sharma, Hundred Strings of Santoor); and six auditory textures, derived from each of these 6 segments (39). To synthesize auditory textures, we used the Auditory Texture Model toolbox (http://mcdermottlab.mit.edu/downloads.html). Textures are sounds that share low-level statistics with the original excerpts from which they are derived. The low-level statistics include average spectral content, amplitude modulation spectra at each frequency channel, as well as correlations between amplitude modulation patterns across frequency channels. Then, starting from white noise tokens, these low-level statistics are imposed on the sound, resulting in new sound segments that share these statistics with the original music segments but are otherwise random. The resulting 12 sounds were high pass filtered (Corner frequency=1500 Hz), as done during the exposure phase. The waveforms were saved as computer files that were used during the experiments (audio files S1-S12). These sounds were presented 3 times each in a pseudo- random sequence, with an ISI of 10 s (2 seconds of silence between successive sound presentations).

### Data analysis

All the analysis was conducted using MATLAB (2020-2022b, The MathWorks).

The duration of each behavior test was divided into nine segments of 10 minutes (habituation) or 20 minutes (music vs. silence test). The proportion of time the mice spent in the two zones in the box in each of these time segments was determined. The habituation test was used to select the less visited zone as the music zone for the test conducted on the next day. The main data of interest were the duration of time spent in the music zone in each of these segments. Since there was only one female mouse in the chirp-exposed group, we analyzed for this group the data from the males only.

The calcium responses analyzed here consist of the average signal over the active pixels, since the spatial dependence of the calcium responses was weak. However, we used the spatial information in order to clean the signal from non-neural signals and in order to identify the active area of the image.

The acquired images were binned from a resolution of 512x512 pixels to 128x128 pixels by averaging blocks of 4x4 pixels. Reduction of motion artifacts was performed in two steps. First, image alignment was performed using the ‘imregister’ function from the MATLAB image processing toolbox using the rigid transformation option. To prevent boundary effects, we excluded the rows and columns at the boundaries, resulting in a 126 x 126 data matrix. The second step used DSS, a denoising algorithm based on spatial filtering (40). The goal was to find spatial patterns that fluctuated periodically at rhythms consistent with heart rate (∼5-10 Hz). These spatial patterns were then subtracted from each image, reducing heart-rate artifacts.

To identify the slowly changing baseline signal of the imaging data, a 3rd degree polynomial was fitted to the temporal signal of each pixel. This polynomial was then subtracted from the raw data, and the mean of the raw data added back (‘aligned data’). Next, the 5th percentile of the aligned data was used as baseline. To estimate ΔF/F (fractional fluorescence change), the baseline was subtracted from the aligned data, and the result was divided by the baseline. The images were then further downsampled to 18 × 18 pixels by taking the mean of every 7×7 pixel block. The final pixel size was 426 × 426 μm^2^.

The processed temporal time course in each pixel was segmented into trials, and the mean response across trials was calculated. To identify active pixels, we used the fact that the corner pixels were expected to exhibit low activity and therefore have relatively low standard deviation over time. The maximum standard deviation among the corner pixels was determined, and a threshold was set to 110% of this value. Pixels exceeding this threshold were categorized as active. This threshold provided a satisfactory definition of the region of active pixels as judged by visual inspection. Finally, the signal was averaged over all active pixels.

The responses to the complex sounds showed clear relationships to the spectro-temporal structure of the sounds (Fig S3A). We therefore analyzed these responses by cross-correlating them with a coarse auditory representation of the sounds. We simulated the peripheral auditory representation using the Bruce auditory nerve model as implemented in the Auditory Nerve Model Toolbox (41, 42). We used 100 frequency channels covering the range of 1000 to 22,000 Hz, using the ‘cat’ option to better fit to the animal data. The resulting fine-grained auditory representation was averaged within 5 non-overlapping 1-octave bands and each band was lowpass-filtered with a corner frequency of 5 Hz, since the calcium signals were not expected to fluctuate at higher rates. The resulting coarse, slowly-varying 5-band auditory representation was then correlated with the recorded calcium signals (Fig. S4 shows an example). This analysis is premised on the hypothesis that the firing rates of the neuronal population that generates the calcium responses depend on the overall excitation reaching auditory cortex, and that excitation in its turn follows largely the amplitude fluctuations of the auditory signal within the relevant frequency bands. The significant across-frequency correlations that were present in the complex sounds used here justified the use of a coarse frequency resolution for this analysis. The duration of the sounds was not sufficiently long to extract spectro-temporal receptive fields from these recordings. Therefore, we only study here the peak correlation values and corresponding latencies, in relationship to the experimental group and sex.

### Statistical analysis

For the statistical analyses, we mostly used linear mixed effects (LME) models, where the fixed effects were group, sex and their interaction, and the random effect consisted of the stimulus identity and mouse identity within group X sex. The significance of the fixed effects was determined using permutation test (n = 1000).Each F-statistic from the ANOVA of the true-data LME model was compared to the distribution of corresponding F-statistics of the appropriately permuted datasets. For testing group effects, we permuted animals across groups, keeping their sex. For testing sex effects, we permuted mouse sex, keeping their group. In consequence we had two estimates for the significance of the sex X group interaction, and we report always the more significant one.

## Acknowledgments

We thank Dr. Dina Moshitch for technical assistance in wide-field calcium imaging.

## Author contributions

Conceptualization: Israel Nelken, Kamini Sehrawat

Methodology: Israel Nelken, Kamini Sehrawat

Investigation: Israel Nelken, Kamini Sehrawat

Visualization: Israel Nelken, Kamini Sehrawat

Funding acquisition: Israel Nelken

Project administration: Israel Nelken

Supervision: Israel Nelken

Writing – original draft: Israel Nelken, Kamini Sehrawat

Writing – review & editing: Israel Nelken, Kamini Sehrawat

## Data and materials availability

All data are available in the main text or the supplementary materials

---

**Fig. S1.**
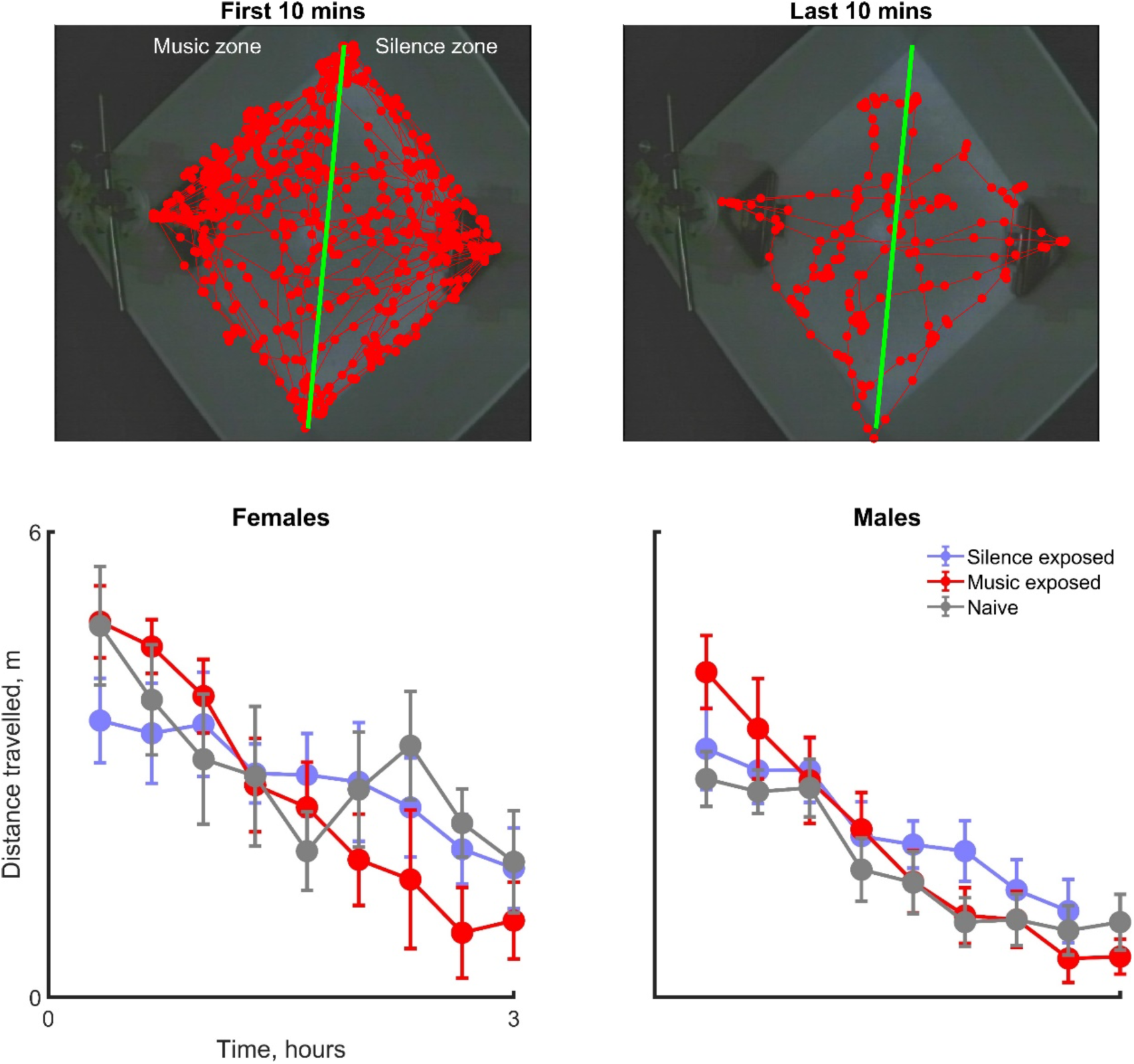
Gradual decrease in exploration during music preference test. (Top) A representative trajectory of a mouse during the first and the last 10 minutes of the music versus silence test (total duration of test: 3 hours), illustrating a gradual decrease in exploration as a function of time. **(Bottom)** Distance travelled during music preference test (x-axis) versus the time of the test (left: females; naïve, n=6; music-exposed, n=6; silence-exposed, n=8; and right: males; naïve, n=10; music-exposed, n=11; silence-exposed, n=10). Statistical summary in Table S1.

**Fig. S2.**
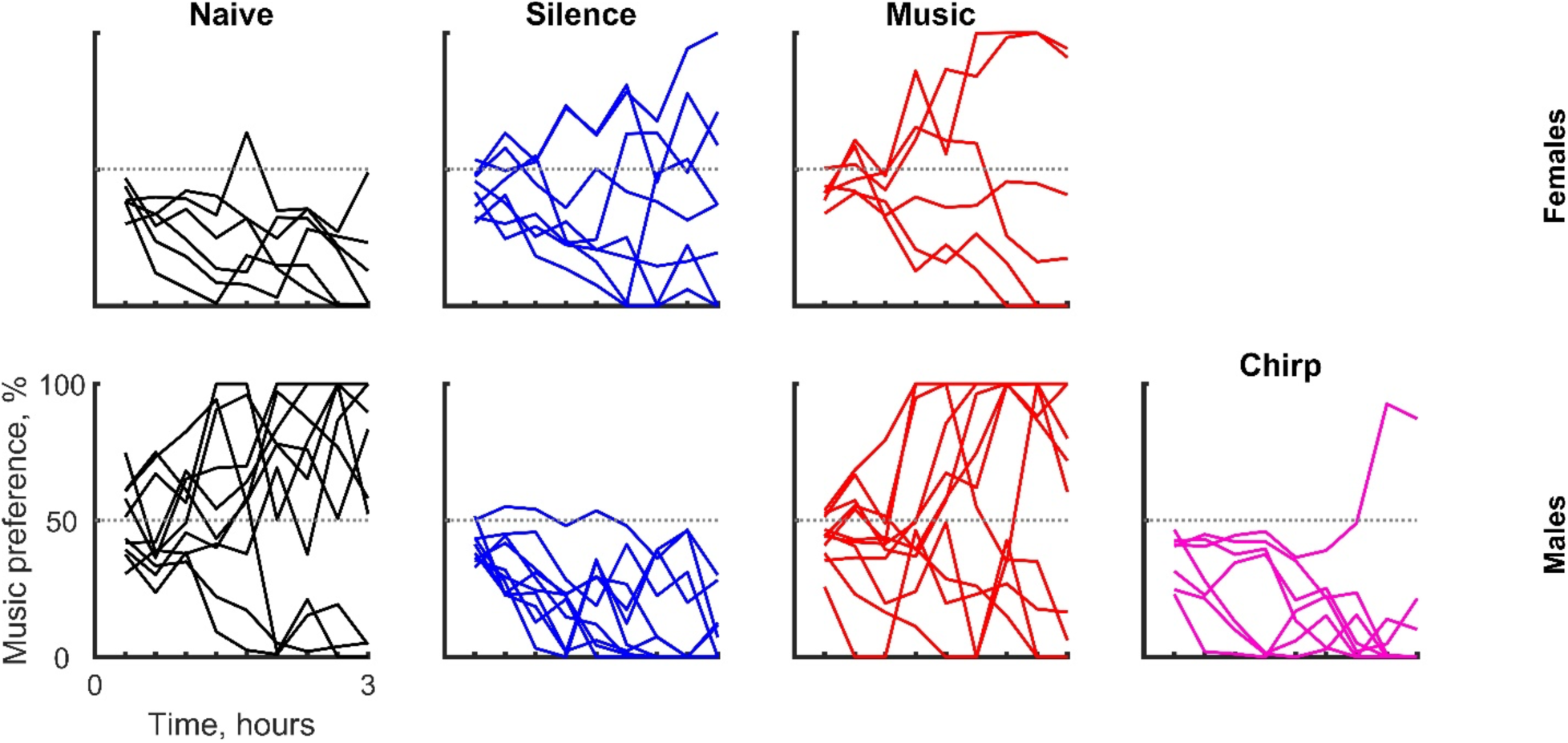
Music preference in individual mice. The fraction of time spent in the music zone (y axis) as a function of time bin (x axis) for each mouse individually. **(Top)** Females (statistical summary in Table S4). **(Bottom)** Males (statistical summary in Table S3).

**Fig. S3.**
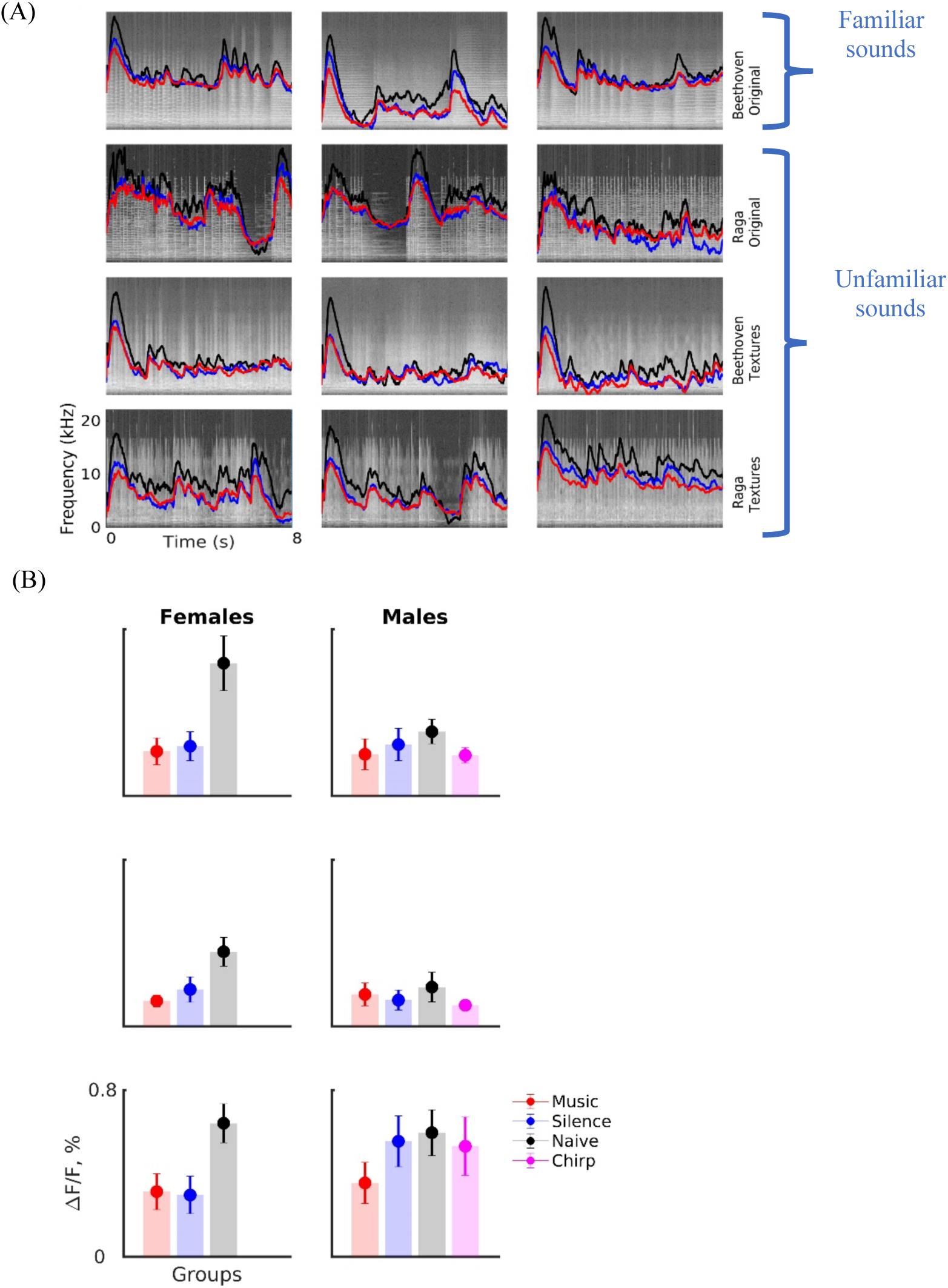
Suppression of neural activity. (A) Average calcium responses (mean ± SE; y axis) as a function of time (x axis) for Beethoven original excerpts or familiar sounds in top row, for Indian raga original excerpts in second row, for textures derived from familiar sounds (see method for details) in third row, and textures of Indian ragas in last row**. (B)** Peak responses (mean ± SE) for different stimuli: Broadband noise (top), Pure tones (middle), and familiar complex sounds (bottom).The response to in chirp-exposed (magenta) male mice exhibited similarities to those in the silence-exposed (blue) male mice but differed significantly from those in the music-exposed males (red; see Table S10).

**Fig. S4.**
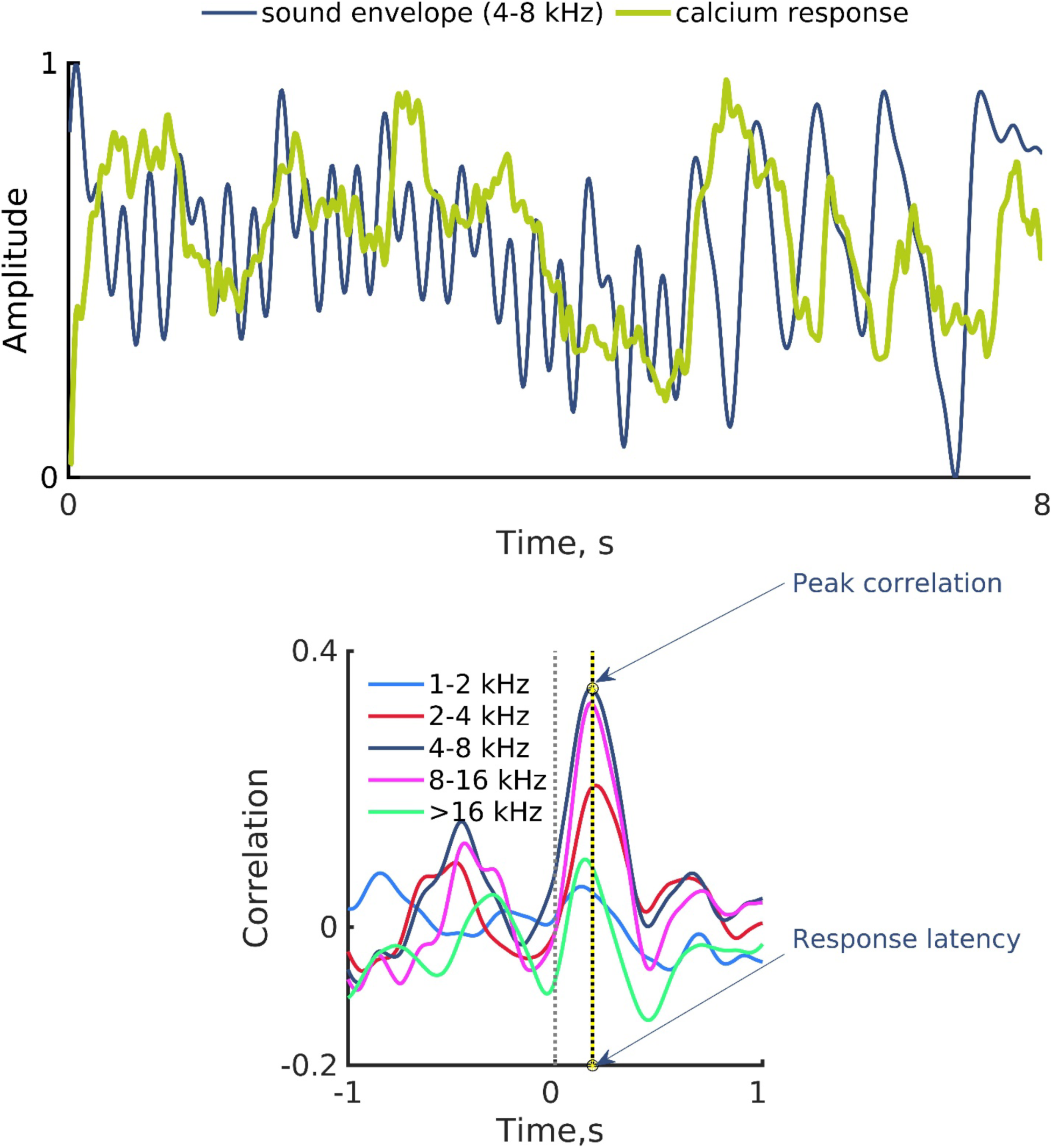
Example of cross-correlation between neural response and band passed sound envelope. (Top) An example of band pass sound envelope (sound Beethoven symphony excerpt 1, 4-8 kHz) and the calcium response evoked by this sound in auditory cortex. **(Bottom)** Cross correlations for five frequency bands of the example depicted in top panel. The peak correlation and corresponding latency for the 4-8 kHz band is marked in yellow.

**Fig. S5.**
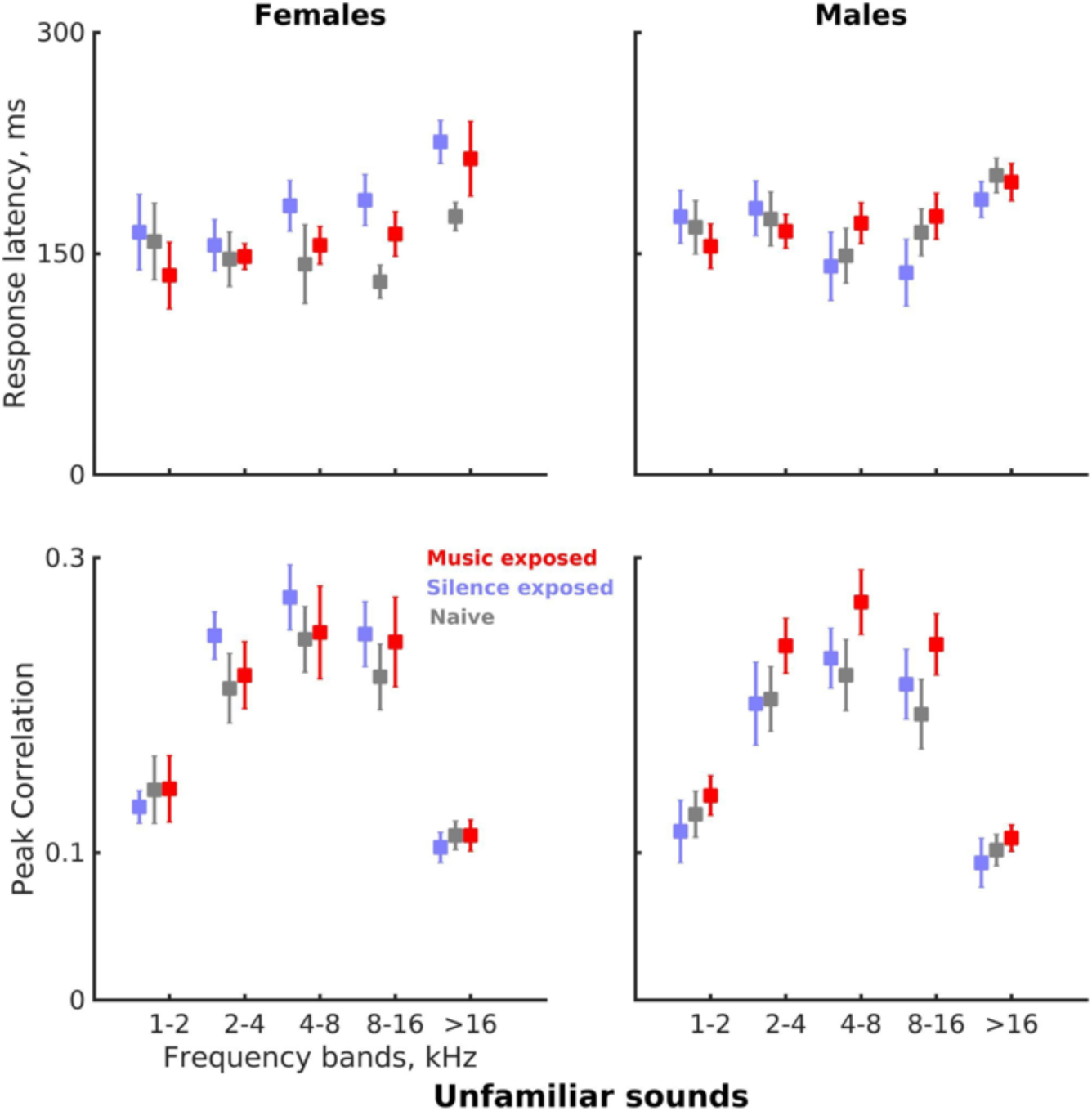
Cross-correlation between neural response and band passed sound envelope of unfamiliar sounds. Same format as Fig. 2C, showing the data for the unfamiliar sounds (see Tables S13 & S14 for the statistical summaries).

**Fig. S6.**
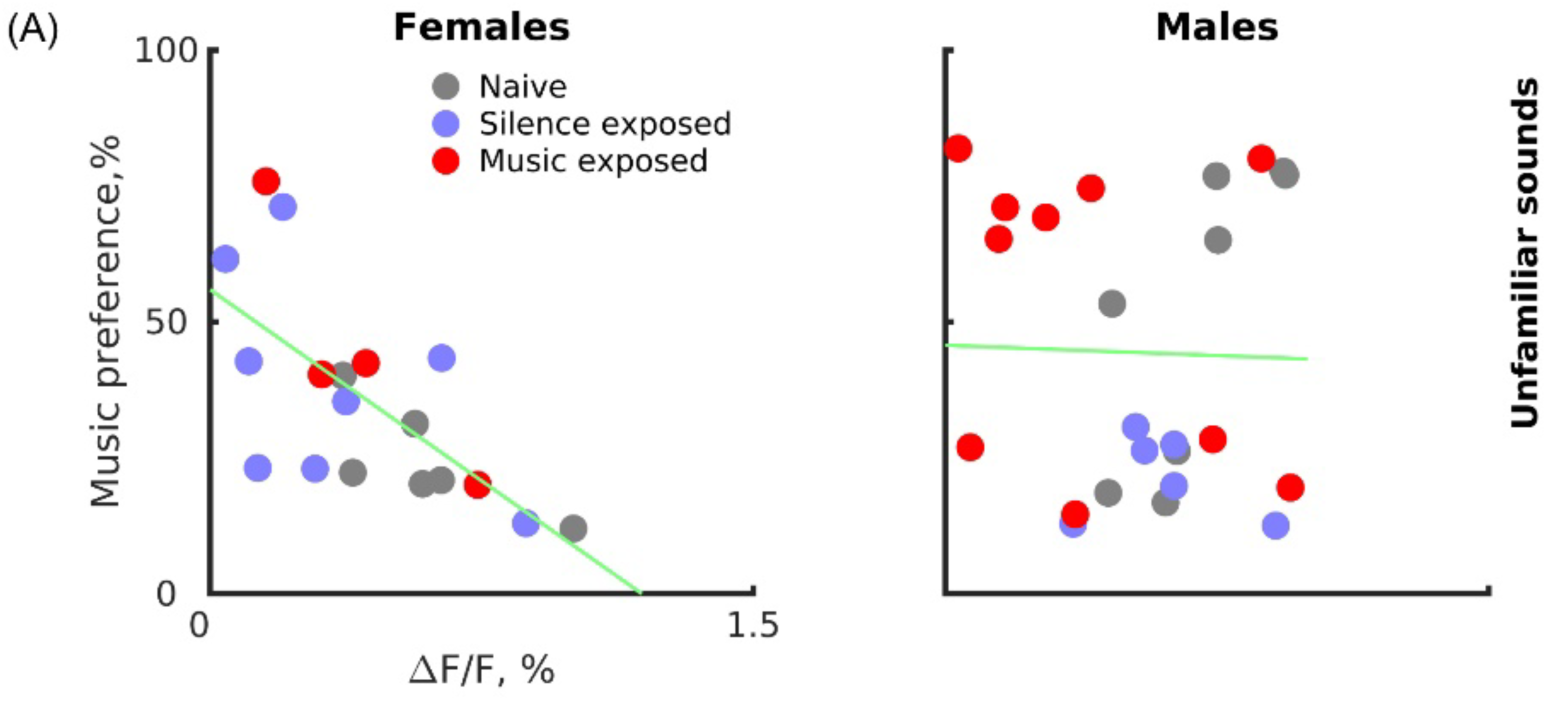
: Neural correlates of behavior. Same display as Fig. 3, for the unfamiliar sounds (see table S17A for statistical summary).

**Table S1.** Statistics for decrease in exploration Linear mixed effects (LME) model for the distance travelled during the test. The fixed effects were experimental groups (naïve: n=16 (M=10, F=6); music-exposed: n=17 (M=11, F=6); and silence-exposed: n=18 (M=10, F=8)), sex, and time in test (9 bins, bin-width=20 mins), with random intercepts for mouse within group X sex. Each F-statistic for the true-data LME model was compared to the distribution of the corresponding F-statistics generated through a permutation test (n=1000). Permutation test was performed for each fixed effect parameter separately. For the interaction term the lowest Pvalue was chosen.

| Term | P |
| --- | --- |
| time in test | <0.001 |
| group | 0.015 |
| sex | <0.001 |
| time in test: group | <0.001 |

**Table S2.** Statistics for the music preference test Linear mixed-effects (LME) model for dwell times in the music zone. The fixed factors were experimental groups, sex (naïve: n=16 (M=10, F=6), music-exposed: n=17 (M=11, F=6)), and silence-exposed: n=18 (M=10, F=8)), time in test (9 bins of 20 min) and their interactions, with random intercepts for mouse within group X sex. Each F-statistic for the true-data LME model was compared to the distribution of the corresponding F-statistics generated through a permutation test (n=1000). Permutation test was performed for each fixed effect parameter separately. For the interaction term the lowest Pvalue was chosen.

| Term | P |
| --- | --- |
| group | 0.016 |
| sex | 0.021 |
| time in test | 0.72 |
| group:sex | 0.007 |
| group:time in test | 0.010 |
| sex:time in test | 0.58 |
| group:sex:time in test | 0.007 |

**Table S3.** Statistics for music preference test for males only. Because of the significant sex X group interaction in Table S2, we fitted sex-specific linear mixed-effects (LME) model to the dwell times in the music zone. The fixed factors were experimental groups (naïve: n=10, music-exposed: n=11, silence-exposed: n=10), time in test and their interactions, with random intercepts for mouse within group. Each F-statistic from the ANOVA of the true-data LME model was compared to the distribution of the corresponding F-statistics generated through a permutation test (n=1000). Permutation test was performed for each fixed effect parameter separately. For the interaction term the lowest Pvalue was chosen.

| Term | P |
| --- | --- |
| group | <0.001 |
| time in test | 0.66 |
| group:time in test | <0.001 |

**Table S4.** Statistics for music preference test for females only. Same as Table S3, for females only (naïve: n=6, music-exposed: n=6, silence-exposed: n=8).

| Term | P |
| --- | --- |
| group | 0.023 |
| time in test | 0.704 |
| group:time in test | 0.606 |

**Table S5.** Statistics for specificity of music exposure for males only. To investigate the specificity of music exposure in influencing the observed preference, we used male mice exposed to chirp sounds (see methods for details). To evaluate the significance of the observed preferences (Fig. S2, bottom) the behavior of the male mice was analyzed using a linear mixed-effects (LME) model similar to the used in Table S3, except that the experimental groups included the additional group of chirp-exposed male mice. The fixed factors were experimental groups (naïve: n=10; music-exposed: n=11; silence-exposed: n=10; chirp-exposed male mice: n=7), time in test (9 bins, 20 mins bin-width) and their interactions, with random intercepts for mouse within group. Each F-statistic from the ANOVA of the true-data LME model was compared to the distribution of corresponding F-statistics generated from (n=1000) permutations. Permutation test was performed for each fixed effect parameter separately. For the interaction term the lowest Pvalue was chosen. There was a main effect of group as well as a highly significant interaction with time in test. To demonstrate that the chirp-exposed and music-exposed mice showed significantly different behaviors, we compared two models: in Model 1 the music-exposed and chirp-exposed animals were grouped together as “sound exposed,” while Model 2 was the previous model in which the two groups were analyzed separately. The statistical significance of the difference in goodness of fit of the two models was estimated using permutation test (n=1000).

|  | Term | P |
| --- | --- | --- |
| Liner mixed effects (LME) model<br>Fixed effects: groups, time in test<br>Random effects: mouse within group<br>(groups: music, silence, chirp, naïve) | group | 0.002 |
|  | time in test | 0.88 |
|  | group:time in test | <0.001 |
| Comparing goodness of fit of two<br>LME models | Model 1<br>(groups: naïve, silence, music, chirp)<br>vs<br>Model 2<br>(groups: naïve, silence, music+chirp) | <0.001 |

**Table S6.**
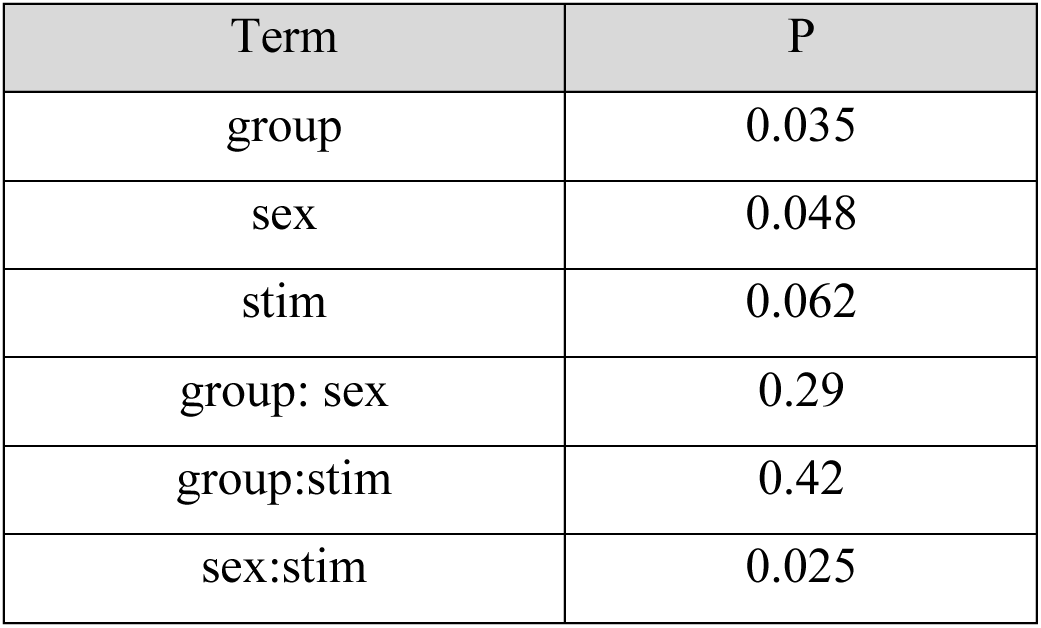
Statistics for suppression in neural responses of exposed groups. Linear mixed-effects model for the neural responses as a function of group and sex. The fixed factors were group, sex (naïve: n=16 (M=10, F=6), music-exposed: n=15 (M=11, F=4)), and silence-exposed: n=15 (M=7, F=8)), and stimulus type (broadband noise, pure tones, familiar sounds), with random intercepts for mouse within group X sex. Each F-statistic from the ANOVA of the true-data LME model was compared to the distribution of corresponding F-statistics generated through a permutation test (n=1000). Permutation test was performed for each fixed effect parameter separately. For the interaction term the lowest Pvalue was chosen.

**Table S7.** Statistics for peak response to familiar complex sounds. Our experimental design included a specific comparison between the silence- and the music-exposed groups (Fig. 2B, bottom). We compared the differences between these peak responses in the two experimental groups (music-exposed: n=15 (M=11, F=4), and silence-exposed: n=15 (M=7, F=8)) using a linear mixed-effects model. The fixed factors were group and sex, with random intercepts for the identity of the stimulus as well as mouse within group X sex. Each F-statistic from the ANOVA of the true-data LME model was compared to the distribution of corresponding F-statistics generated through a permutation test (n=1000). Permutation test was performed for each fixed effect parameter separately. For the interaction term the lowest Pvalue was chosen.

| Term | P |
| --- | --- |
| group | 0.002 |
| sex | 0.34 |
| group: sex | 0.029 |

**Table S8.** Statistics for peak response to pure tones. Responses to pure tones in the music- and silence-exposed mice (Fig. 2B, middle). Same test as Table S7 (music-exposed: n=15 (M=11, F=4), and silence-exposed: n=15 (M=7, F=8)). Each F-statistic from the ANOVA of the true-data LME model was compared to the distribution of corresponding F-statistics generated through a permutation test (n=1000). Permutation test was performed for each fixed effect parameter separately. For the interaction term the lowest Pvalue was chosen.

| Term | P |
| --- | --- |
| group | 0.036 |
| sex | 0.13 |
| group:sex | 0.003 |

**Table S9.** Statistics for peak response to broadband noise. Responses to broadband noise in the music- and silence-exposed mice (Fig. 2B, top). Same test as Table S7 (music-exposed: n=15 (M=11, F=4), and silence-exposed: n=15 (M=7, F=8)). Each F-statistic from the ANOVA of the true- data LME model was compared to the distribution of corresponding F-statistics generated through a permutation test (n=1000). Permutation test was performed for each fixed effect parameter separately. For the interaction term the lowest Pvalue was chosen.

| Term | P |
| --- | --- |
| group | 0.84 |
| sex | 0.94 |
| group: sex | 0.88 |

**Table S10.**
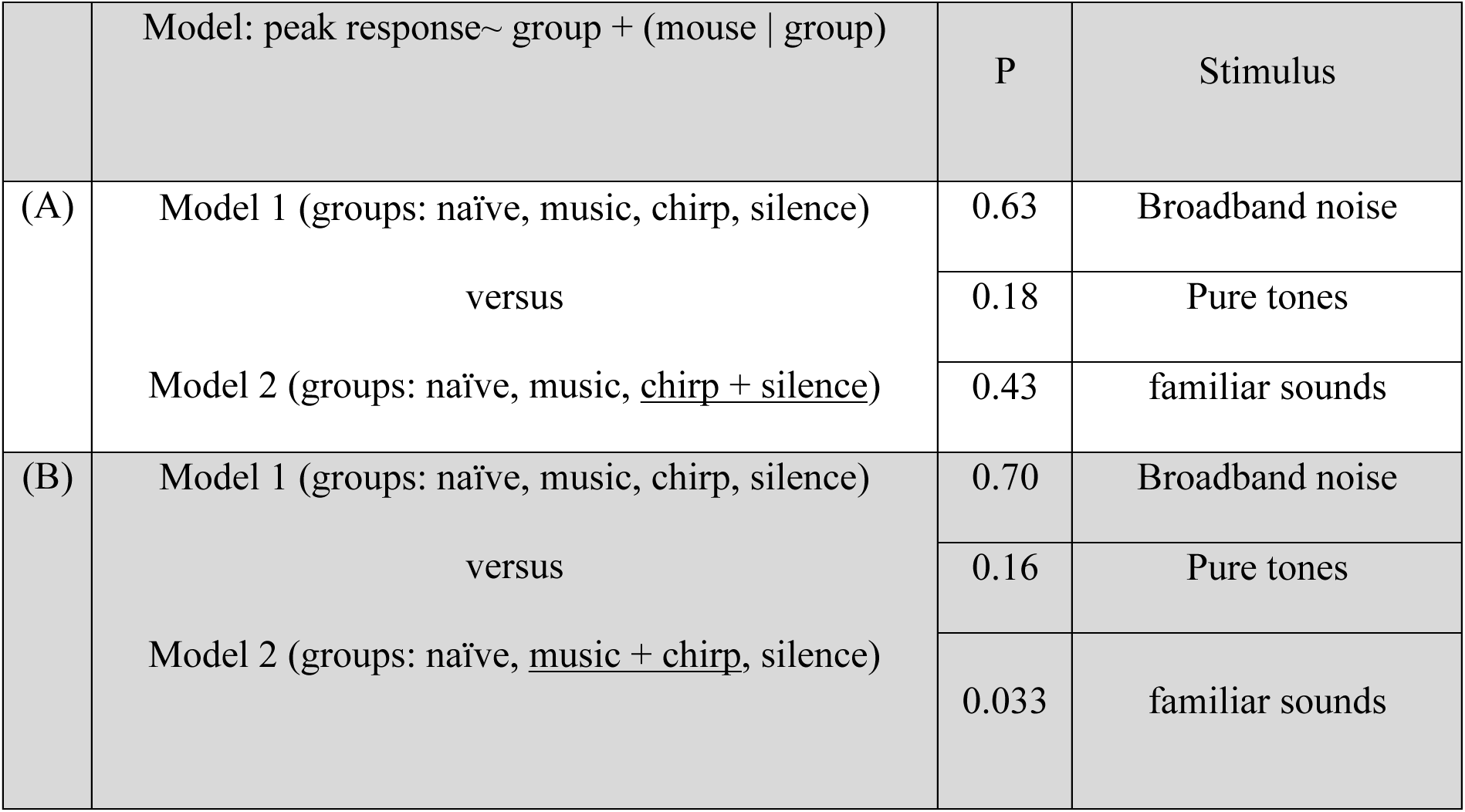
Statistics for similarities in the peak responses of chirp-exposed with silence- exposed and difference from music-exposed male mice. The peak responses of chirp exposed male mice (Fig. S3B; magenta) showed resemblance with silence-exposed males (blue) but different from that of the music-exposed males (red). To demonstrate that the peak neural responses of chirp-exposed male mice were similar to silence-exposed mice (see below A) and different from music-exposed male mice (see below B), we compared goodness of fit of two models for each observation. In both models, the peak neural responses were modelled with the fixed effect of group and random intercept of mouse within group. Significance was verified using permutation test (n=1000). **(A)** One model, Model 2, grouped together the silence- exposed and chirp-exposed groups, while the other model (Model 1) separated them. Non-significant P values indicate that these models were not statistically different from one another, suggesting that chirp-exposed and silence-exposed shared similarities in neural responses. **(B)** One model, Model 2, combined the music-exposed and chirp-exposed groups, while the other model (Model 1) separated them. The difference between music-exposed and chirp exposed male mice was more pronounced in the peak responses to familiar sounds (P=0.033).

**Table S11.**
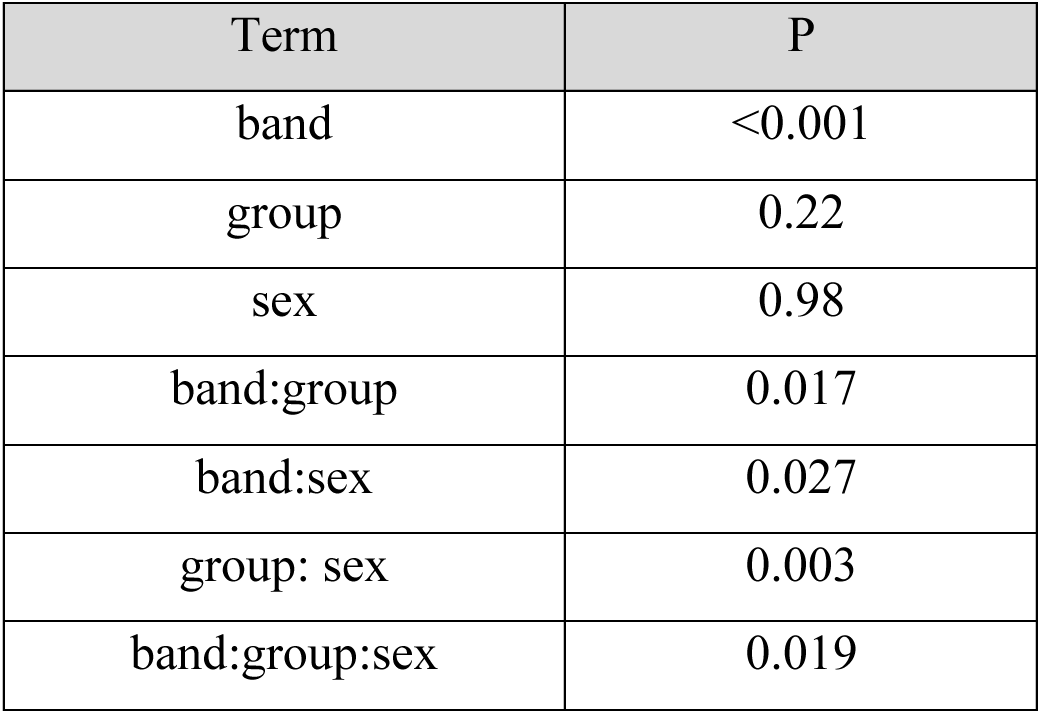
Statistics for response latencies for familiar sounds. We correlated the neuronal calcium signal with the simulated peripheral auditory representation of the familiar sounds in 1-octave bands (see methods for details and Fig. S4). The estimated response latencies (see Fig. 2C, top) defined as the time of peak correlation, were used to verify the relationships between the calcium signal and spectro- temporal structure of the sounds. We used a Linear mixed-effect models with fixed effects of group (naïve: n=16 (M=10, F=6), music-exposed: n=15 (M=11, F=4)), and silence-exposed: n=15 (M=7, F=8)), sex, and frequency band (band1:1-2 kHz, band2: 2-4 kHz, band3: 4-8 kHz, band4: 8-16 kHz, band5: >16 kHz), and random effects of mouse within group X sex. Each F-statistic from the ANOVA of the true-data LME model was compared to the distribution of corresponding F-statistics generated through a permutation test (n=1000). Permutation test was performed for each fixed effect parameter separately. For the interaction term the lowest Pvalue was chosen.

| Term | P |
| --- | --- |
| band | <0.001 |
| group | 0.22 |
| sex | 0.98 |
| band:group | 0.017 |
| band:sex | 0.027 |
| group: sex | 0.003 |
| band:group:sex | 0.019 |

**Table S12.**
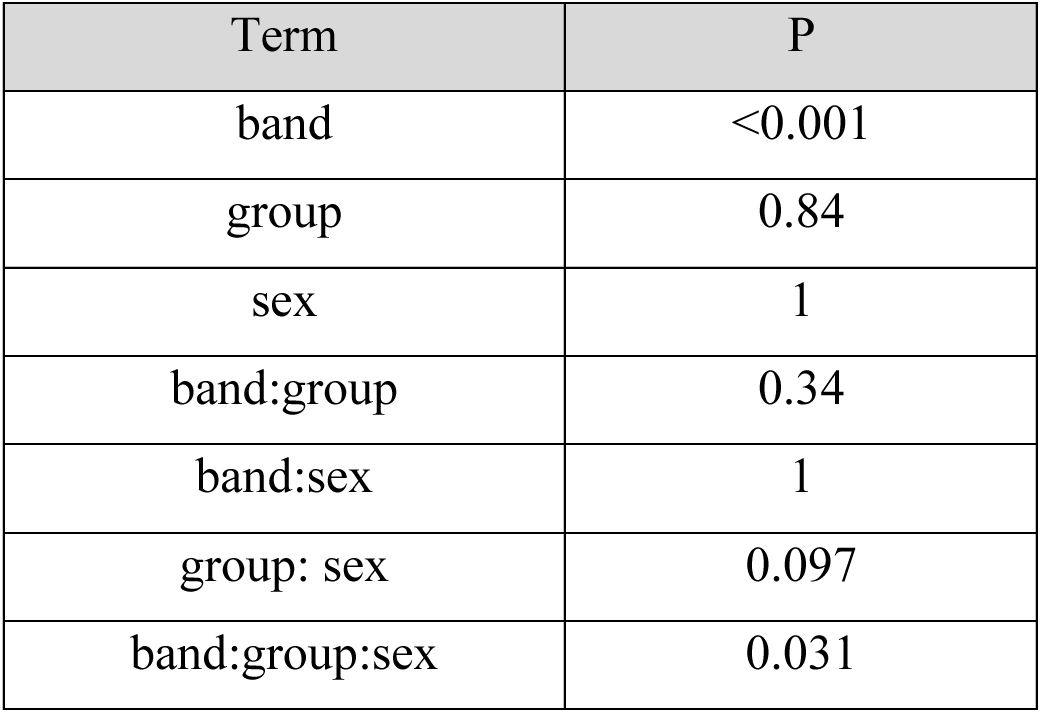
Statistics for peak correlations for familiar sounds. We tested the significance of peak correlations between the neuronal calcium signal with the simulated peripheral auditory representation of the familiar sounds in 1-octave bands (Fig. 2C, bottom) using a Linear mixed-effect models same as Table S11. Each F-statistic from the ANOVA of the true-data LME model was compared to the distribution of corresponding F-statistics generated through a permutation test (n=1000). Permutation test was performed for each fixed effect parameter separately. For the interaction term the lowest Pvalue was chosen.

**Table S13.** Statistics for response latencies for unfamiliar sounds. The latencies of peak correlations between the neuronal calcium signal with the simulated peripheral auditory representation of the unfamiliar sounds in 1-octave bands (see Fig. S5, top) was tested for statistical significance using a Linear mixed-effects model same as Table S11. Each F-statistic from the ANOVA of the true-data LME model was compared to the distribution of corresponding F-statistics generated through a permutation test (n=1000). Permutation test was performed for each fixed effect parameter separately. For the interaction term the lowest Pvalue was chosen.

| Term | P |
| --- | --- |
| band | <0.001 |
| group | 0.34 |
| sex | 0.42 |
| band:group | 0.57 |
| band:sex | 0.039 |
| group: sex | 0.35 |
| band:group:sex | 0.16 |

**Table S14.** Statistics for peak correlations for unfamiliar sounds. The peak correlations between the neuronal calcium signal with the simulated peripheral auditory representation of the unfamiliar sounds in 1-octave bands (see Fig. S5, bottom) was tested for statistical significance using a Linear mixed-effects model same as Table S11. Each F-statistic from the ANOVA of the true-data LME model was compared to the distribution of corresponding F-statistics generated through a permutation test (n=1000). Permutation test was performed for each fixed effect parameter separately. For the interaction term the lowest Pvalue was chosen.

| Term | P |
| --- | --- |
| band | <0.001 |
| group | 0.58 |
| sex | 0.26 |
| band:group | <0.001 |
| band:sex | 0.33 |
| group: sex | 0.020 |
| band:group:sex | 0.62 |

**Table S15.** Statistics for neural activity and behavior. The relationship between animal behavior and the peak calcium responses, is shown in Fig. 3 and Fig. S6. To assess the statistical significance of this relationship, we used linear mixed-effect models for each stimulus type separately (BBN, pure tones, and complex sounds; all sounds together (Fig. 3), familiar sounds (Fig. 3) and unfamiliar sounds (Fig. S6). The response variable was overall time spent in music zone during the Music vs Silence Test. The fixed effects were group (naïve: n=16 (M=10, F=6), music-exposed: n=15 (M=11, F=4)), and silence-exposed: n=15 (M=7, F=8)), sex, and peak calcium responses. And the random intercepts for mouse within group X sex). Significance was assessed using permutation tests (n=1000). Permutation test was performed for each fixed effect parameter separately. For the interaction term the lowest Pvalue was chosen.

|  | Term | P | Stimulus |
| --- | --- | --- | --- |
| (A) | group | 0.006 | Broadband noise<br>(BBN) |
|  | sex | 0.505 |  |
|  | neural response | 0.17 |  |
|  | group:sex | 0.018 |  |
|  | group: neural response | 0.048 |  |
|  | sex:neural response | 0.042 |  |
| (B) | group | 0.01 | Pure tones |
|  | sex | 0.13 |  |
|  | neural response | 0.22 |  |
|  | group:sex | 0.005 |  |
|  | group: neural response | 0.053 |  |
|  | sex:neural response | 0.30 |  |
| (C) | group | 0.026 | Complex sounds<br>(All) |
|  | sex | 0.94 |  |
|  | neural response | 0.43 |  |
|  | group:sex | 0.042 |  |
|  | group: neural response | 0.22 |  |
|  | sex:neural response | 0.049 |  |
| (D) | group | 0.051 | Familiar sounds |
|  | sex | 0.89 |  |
|  | neural response | 0.44 |  |
|  | group:sex | 0.047 |  |
|  | group: neural response | 0.26 |  |
|  | sex:neural response | 0.16 |  |
| (E) | group | 0.022 | unfamiliar sounds |
|  | sex | 0.94 |  |
|  | neural response | 0.45 |  |
|  | group:sex | 0.034 |  |
|  | group: neural response | 0.24 |  |
|  | sex:neural response | 0.039 |  |

**Table S16.** Dependence of time in music zone and neural responses, by sex. Since there were significant interactions between group and sex for all sound types (Table S15), we further investigated the relationship between neural activity and behavior for males and females separately. The peak responses to different stimuli were modelled separately using linear mixed effects model. We selected the best fit model by comparing the goodness of fit of two models. In model 1, the fixed effects were groups, and behavior response along with random intercepts of mouse within group. In model 2, the fixed effect was only behavior response. For all stimuli, these two models were not statistically different, both in females (BBN: P=0.12; pure tones: P=0.22; Complex sounds: P=0.507) and in males (BBN: P=0.20; pure tones: P=0.33; Complex sounds: P=0.403). Therefore, we continued analyzing the relationship between neural activity and behavior for males and females separately, using the linear mixed effects model with the fixed effect of behavior response only. Each F-statistic from the ANOVA of the true-data LME model was compared to the distribution of corresponding F-statistics generated through the permutation process (n=1000). Permutation test was performed for each fixed effect parameter separately. For the interaction term the lowest Pvalue was chosen.

| Stimulus | males | females |
| --- | --- | --- |
| Broadband noise (BBN) | 0.48 | 0.030 |
| Pure tones | 0.91 | 0.040 |
| Complex sounds (All) | 0.82 | 0.002 |
| Familiar sounds | 0.62 | 0.006 |
| Unfamiliar sounds | 0.95 | 0.002 |

**Table S17.** Correlations between behavior response and neural activity, by sex and stimulus type. For a simplified interpretation of the results shown in Table S15 and Fig. 3, we estimated the correlation coefficients using MATLAB function “coefftest” between neural response and behavior for males (naïve: n=10, music-exposed: n=11 males, and silence-exposed: n=7) and females (naïve: n=6, music-exposed: n=4, and silence-exposed: n=8 females) separately. The R statistic from the “coefftest” was compared to the distribution of statistics of “coefftest” generated from n=1000 permutations.

| Stimulus | males | females |
| --- | --- | --- |
| Broadband noise (BBN) | 0.47 | 0.030 |
| Pure tones | 0.89 | 0.028 |
| Complex sounds (All) | 0.82 | 0.002 |
| Familiar sounds | 0.62 | 0.004 |
| Unfamiliar sounds | 0.87 | 0.001 |

